# Navigating the Night: Effects of Artificial Light on the Behaviour of Atlantic Puffin Fledglings

**DOI:** 10.1101/2024.04.24.590944

**Authors:** T.M. Brown, S.I. Wilhelm, A.D. Slepkov, K. Baker, G.F. Mastromonaco, G. Burness

**Affiliations:** Environmental and Life Sciences Graduate Program, Trent University; Environment and Climate Change Canada; Department of Physics & Astronomy, Trent University; Department of Biology, Trent University; Wildlife Science Division, Toronto Zoo

**Keywords:** ALAN, alcid, behaviour, fledging, light attraction, phototaxis, seabird

## Abstract

Every year in Newfoundland, young Atlantic Puffins (*Fratercula arctica*) departing their nests at night for the first time become stranded in towns near their breeding colonies, a phenomenon thought to be caused by attraction toward artificial light. To test this hypothesis, we conducted three behavioural experiments. First, we illuminated beaches near a breeding colony to determine whether more fledglings would become stranded in illuminated versus dark conditions. Next, we conducted a Y-maze experiment to test stranded fledglings for phototactic behaviour in general and for preferences among High Pressure Sodium (HPS), Warm White Light-Emitting Diode (LED), Cool White LED, Blue LED, and Orange LED light. Lastly, we quantified activity levels of stranded fledglings in an open field test during exposure to several different light types. We found significantly more fledglings stranded when beaches were illuminated, and fledglings significantly preferred light over darkness in the Y-maze, supporting our hypothesis that Atlantic Puffin fledglings become stranded due to light attraction. Fledglings displayed no preferences for certain light types over others in the Y-maze, potentially suggesting that strandings in this species may not be mitigable by changing the streetlight type in stranding-prone towns. Interestingly, fledglings exhibited higher activity levels in darkness and HPS light than in LED light, potentially holding implications for rescue, rehabilitation, and husbandry programs. Overall, our findings demonstrate that the only evidence-based strategy for the reduction of Atlantic Puffin strandings remains to be the reduction of coastal artificial lighting; however, further research is needed to determine if aspects of artificial light besides bulb type may be altered to effectively reduce light attraction in this species.

## INTRODUCTION

As global human development continues to increase, artificial light at night (ALAN) is of growing concern as a pollutant of dark nightscapes (Jägerbrand and Spoelstra, 2023). In addition, the widespread conversion to energy-efficient Light-Emitting Diode (LED) lighting is expected to worsen the ecological impacts of ALAN through a variety of mechanisms (Pawson and Bader, 2014; Davies and Smyth, 2018). These mechanisms are both physiological (e.g., many organisms are more sensitive to short-wavelength light, which is prevalent in LED outputs, than to the long-wavelength light that dominates light types used previously, such as sodium vapour) and logistical (e.g., greater energy-efficiency allows for increased light production at the same or lower energy cost as other lighting technologies; Pawson and Bader, 2014; Davies and Smyth, 2018) in nature. Across taxa, disruption of natural diel light cycles by the presence of ALAN can dysregulate circadian rhythms and lead to endocrinological dysfunction, behavioural changes, and decreased fitness, among other effects (Sanders et al., 2021; Yang et al., 2024). One way in which ALAN can alter behaviour, especially that of nocturnal animals, is by acting as a supernormal visual stimulus that induces phototactic behaviour (either positive or negative; Rich and Longcore, 2006). Positive phototaxis (or photopositivity), which is often ascribed to light attraction, has been documented in various animal taxa including insects, amphibians, fish, sea turtles, migratory songbirds, and seabirds, among others (Rich and Longcore, 2006; Rodríguez et al., 2017a; Jägerbrand and Spoelstra, 2023; Fabian et al., 2024). Positive phototaxis toward sources of ALAN can result in mortality due to collisions with glass windows, lighted structures, and vehicles; stranding and dehydration or desiccation in unfamiliar locations; and predation of stranded individuals following disorientation and/or collision (Witherington, 1997; Rich and Longcore, 2006; Rodríguez et al., 2017a; Van Doren et al., 2021).

Seabirds of the orders Procellariiformes and Charadriiformes (family Alcidae) are prone to becoming stranded in human settlements following phototaxis toward and disorientation by ALAN (Rodríguez et al., 2017a; but there are other non-mutually exclusive hypotheses for stranding such as simple navigation error during natural dispersal; Brown et al., 2023). Fledglings of burrow-nesting and night-fledging seabird species appear to be the most susceptible to light-induced stranding, although adults of some species are also affected (Rodríguez et al., 2017a; Burt et al., 2024). There is mounting evidence that suppression of ALAN near colonies of burrow-nesting, night-fledging procellariiform seabirds can reduce the number of stranded fledglings (Reed et al., 1985; Miles et al., 2010; Rodríguez et al., 2014; Burt et al., 2024). However, it is not always possible to eliminate or suppress the use of ALAN, especially at industrial and commercial sites where lighting is necessary for regular operation and human safety. There is thus a growing interest from industry, non-governmental conservation organizations, and various levels of government in investigating and implementing alternate “less attractive” light types where suppression or elimination of ALAN is not possible (e.g., Mercer, 2018). Limited evidence suggests that lights with high correlated colour temperature (CCT; a relatively crude but simple and widely-used metric of the perceived colour of a nominal white light source; Durmus, 2022) and predominantly blue-violet wavelengths (such as metal halide types) may be more attractive to fledgling procellariiforms and other wildlife than those with low CCT and red-orange dominant spectra (such as sodium vapour types; Salamolard et al., 2001 as cited in Minatchy, 2004; Rodríguez et al., 2017b; Longcore et al., 2018), but other findings suggest the opposite (Atchoi et al., 2024) and some indicate no difference (Urmston et al., 2022). An increased attraction toward blue-violet wavelengths may be explained by an increased sensitivity to these wavelengths by the visual systems of the affected taxa (Reed, 1986; Bowmaker et al., 1997; Hart, 2004; Pawson and Bader, 2014; Davies and Smyth, 2018).

While research interest in procellariiform light attraction has increased in recent years, comparatively little attention has been given to other seabirds that are similarly affected by ALAN. Most notably, the Atlantic Puffin (*Fratercula arctica*) represents the only non-procellariiform seabird species that gets stranded routinely in coastal communities, with insular Newfoundland (Newfoundland and Labrador, Canada) and Iceland being the best-known affected locations (Rodríguez et al., 2017a; Wilhelm et al., 2021; Katz, 2023). Adults are diurnal, but fledglings leave their burrows on maiden flights to the sea at night, unattended by parents, leaving them susceptible to the influence of ALAN. Although populations in the western Atlantic remain robust and are even increasing (Wilhelm et al., 2015), those in the eastern Atlantic are declining, likely due to low recruitment (Miles et al., 2015; Fayet et al., 2021). Thus, with increasing coastal development, understanding the extent and spectral attributes through which ALAN impacts post-fledging behaviour in this species is urgently needed.

In Newfoundland, certain well-lit commercial and industrial locations near Atlantic Puffin breeding colonies are known to consistently yield stranded puffin fledglings (hereafter, “pufflings”) and have thus been deemed stranding “hotspots” (Canadian Parks and Wilderness Society – Newfoundland and Labrador Chapter, unpublished data). The existence of such “hotspots” has been taken as anecdotal evidence in support of the light attraction hypothesis of stranding in puffins, although it is unclear to what degree this pattern may be driven by a spatial bias in non-systematic search effort (e.g., toward well-lit areas, where people are also presumably more comfortable searching at night). Furthermore, while hundreds of puffins are stranded annually, this represents only a very small proportion of the total number of individuals fledging from the colonies (Wilhelm et al. 2021), providing support to the natural random dispersal hypothesis which suggest that young birds become stranded because of their inexperience in navigating a new environment (Brown et al., 2023). To date, no experimental study has been conducted to directly test the light attraction hypothesis and confirm the role of ALAN in causing Atlantic Puffin strandings. To do so would require a coarse-scale experimental approach wherein, at a minimum, light levels (including specifically a “dark” condition), geographic location, and search effort are all tightly controlled (Brown et al., 2023).

At small geographic scales, the behavior of pufflings and other stranding-prone seabird fledglings exposed to ALAN remains largely unknown but can be studied ex- or in-situ (e.g., in a Y- or T-maze choice experiment). In the only alcid whose fine-scale phototactic behaviour has been studied to date, Ancient Murrelet (*Synthliboramphus antiquus*) chicks oriented more often toward a reflected light source than toward darkness (Gaston et al., 1988). Stranded Cory’s shearwater (*Calonectris borealis*) fledglings, however, consistently chose darkness over light in a similar experiment (Atchoi et al., 2024). In addition to testing for the presence of photopositivity in general (i.e., comparing taxic responses to light versus dark), it is also of interest whether stranding-prone seabird fledglings exhibit fine-scale preferential phototactic responses or differential activity levels in response to certain spectra of artificial light over others. For example, Atchoi et al. (2024) found that both fledgling and adult Cory’s shearwaters significantly preferred red over blue light when both were provided as options simultaneously. Such fine-scale experiments may not perfectly mimic what occurs in nature, but they nonetheless inform our understanding of behavioural responses to light, especially in ground-based contexts. This information holds implications for rescue efforts in predicting where stranded seabirds are likely to go and how they will behave following stranding, which should allow for higher rescue success.

In this study, we tested the effect of ALAN on puffin behaviour at both a coarse and fine scale. At a coarse scale, we experimentally tested the light attraction hypothesis of stranding in the field by directing a 30 000 lumen LED light toward the Gull Island (Witless Bay) Atlantic Puffin breeding colony, on alternate nights from two different dark beaches during the fledging season. Given the hypothesis that pufflings become stranded by light attraction, we expected that more pufflings would be observed at the beaches when they were illuminated than when the beaches were dark. To test for effects of ALAN on a fine scale, we used a Y-maze to evaluate the propensity of already-stranded pufflings to exhibit phototactic behaviour and preferences for various artificial light spectra. We expected that fledglings would exhibit positive phototaxis in the Y-maze by preferentially choosing light over darkness when both were provided as options. We further expected that when pufflings were exposed to two different artificial light types, they would preferentially choose those with “cooler” hues (e.g., Cool White LED; Blue LED) over those with “warmer” hues (e.g., High Pressure Sodium (HPS); Warm White LED; Orange LED). This reasoning is based on the peak optical sensitivity of closely-related auks to 406 nm violet light and that of stranding-prone procellariiforms to 402-405 nm violet light (Ödeen and Håstad, 2003). To determine if different spectra of artificial light induce different activity levels in pufflings, we conducted an open field test. Although we expected to observe differences in activity levels among the light treatments, particularly when compared with darkness, we could not predict directionality *a priori*.

## METHODS

### Ethical Note

All procedures involving the use of animals for research were performed in accordance with Trent University’s Animal Care Protocol guidelines under the Canadian Council on Animal Care (CCAC), protocols #26223 and #28033, and Canadian Wildlife Service guidelines under scientific permit # SC4051. All experiments were compliant with current Canadian law. Birds were captured using handheld mesh nets, were not manipulated beyond simple handling, and were held in captivity for as short a time as possible. We further attempted to reduce the stress experienced by birds by housing them individually in quiet and dimly lit conditions for the duration of captivity.

### Study Sites and Species

In Newfoundland, most stranded pufflings are found along a ∼15 km section of coastline between the communities of Bay Bulls and Bauline East (Figure 1). This section forms the focal search area for the Canadian Parks and Wilderness Society – Newfoundland and Labrador Chapter’s (CPAWS-NL) “Puffin Patrol”, a volunteer-based puffling rescue program that operates each year during the fledging (and stranding) season (Wilhelm et al., 2013, 2021). The Witless Bay Ecological Reserve, which borders this coastline, hosts close to 400 000 breeding pairs of Atlantic Puffins, forming the largest concentration in North America (Lowther et al., 2020; SIW, unpublished data). Stranded pufflings are thought to largely originate from the two largest breeding colonies on Gull Island (47.262297°N, 52.773389°W, estimated at 168 000 pairs; SIW, unpublished data) and Great Island (47.187182°N, 52.813495°W, estimated at 204 000 pairs; SIW, unpublished data), both part of the reserve and located less than 3 km from shore at their closest points (Figure 1).

**Figure 1.**
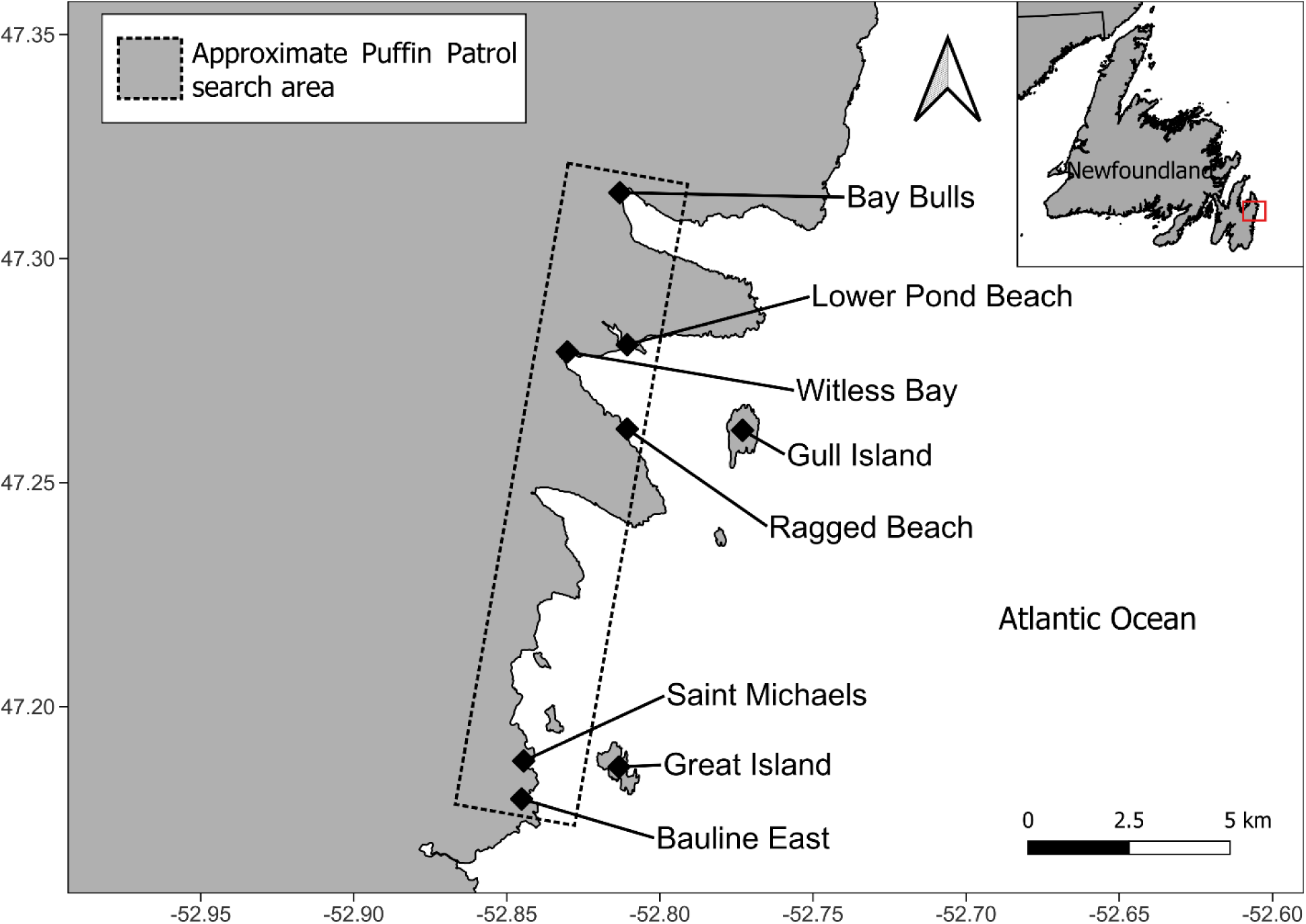
Map of Puffin Patrol search area in Newfoundland, Canada, including all relevant locations of research activities (Bay Bulls, Lower Pond Beach, Ragged Beach, Saint Michaels) and the two largest Atlantic Puffin breeding colonies (Gull Island, Great Island).

To study the behavioural response of pufflings to light at coarse spatial scales, we performed an “illuminated beach experiment” (hereafter, beach experiment) in August 2022 at two sites near Witless Bay, Newfoundland: Lower Pond Beach (47.281464°N, 52.810982°W) and Ragged Beach (47.262068°N, 52.811644°W; Figure 1). Both of these beaches face Gull Island: Lower Pond Beach at a distance of approximately 3.1 km, and Ragged Beach at a distance of approximately 2.5 km.

To study the fine-scale spatial behavioural response of pufflings to lights of differing spectra, we performed both a Y-maze “choice experiment” and an open field test in August 2021 in Bay Bulls, Newfoundland and Labrador, Canada (47.315178°N, 52.813887°W; Figure 1). In 2021, we conducted the fine-scale experiments in one of two identical “field labs” (storage trailers, 2.4 × 1.4 × 2.1 m) located adjacent to the headquarters for CPAWS-NL’s annual Puffin Patrol rescue program (and the designated drop-off location for all rescued pufflings). One “lab” was used for the choice experiment, while the other was for the open field test. In August 2022, we tested one additional choice experiment combination in a building in Saint Michaels (47.187247°N, 52.843990°W; Figure 1).

### Response of Pufflings to Light at a Coarse Spatial Scale

#### Materials

We illuminated one of two beaches (Lower Pond beach or Ragged Beach; Figure 1) over 11 nights during the fledging season between 15-27 August 2022, coinciding with a transition from 85% to 0% lunar illumination (moongiant.com). Experiments were not conducted August 17 and 18 due to inclement weather. At both beaches, a 30 000 lumen, 6390 K LED light apparatus (Willpower 52-inch, 675-Watt LED light bar: Xalapa, Veracruz, Mexico; spectrum displayed in Appendix Figure A1) was mounted to a 1.5-metre wooden fence board and placed on two overturned plastic buckets at the highest point of the beach (approximately 2 metres above sea level) such that the light sat horizontal and parallel to the beach, facing Gull Island. On each night during the experimental period, we illuminated one beach while the other remained dark. At the Light beach, the light apparatus was connected to a Discover 12V 35AH/20HR rechargeable deep cycle marine battery (Richmond, B.C., Canada) via a wiring harness with an in-line fuse and toggle power switch; at the Dark beach the battery and the light apparatus with attached wiring harness were set up but not connected.

#### Experimental procedure

A coin was flipped to determine which of the two beaches was illuminated on the first night of the experiment; thereafter, the Light treatment was alternated between beaches. Because there was one illuminated beach and one unilluminated beach every night, we have *N* = 11 “Light” and *N* = 11 “Dark” experimental trials. Regardless of treatment (Light or Dark), there was a team of usually two to three (one instance of four, and one instance of five) biologists and trained volunteers stationed at each beach to operate the light apparatus and to observe and capture pufflings for the duration of the experiment. Trials ran for exactly two hours, sometime between 2130 and 0030 hours (to coincide with the peak stranding period; CPAWS-NL, unpublished data), depending on the night. On two nights there was an equipment malfunction in the middle of the trial, but by extending the Light treatment trial we ensured the total duration of two hours of light exposure remained the same.

Approximately 10 minutes before the experiment start time, 1-2 team members at each beach conducted a brief survey to a minimum of 25 metres on either side of the light apparatus and scanned the water with the unaided eye and night vision binoculars (Bestguarder NV900 4.5-22.5 × 40 HD; Shenzhen, China) as thoroughly as allowable by the weather conditions, to ensure no pufflings were already present in the vicinity prior to the start of the experiment. We then placed two reflective safety vests 25 metres on either side of the light apparatus and two vests closer to the water’s edge at the same distance (total four vests on each beach) to delineate the focal observation area (50 metres long and varying from 15-20 metres wide = ∼750-1000 m^2^). This distance was chosen based on preliminary tests of the resolution and range of the night vision binoculars and the width of the light beam. There was no way to consistently delineate the focal observation area over water, but the best effort was made to estimate the relative distances of pufflings observed.

During the two-hour experiment period on the Light and Dark beaches, observations principally consisted of one team member at each beach visually scanning the focal beach observation area with night vision binoculars; however, it should be noted that on most nights, even the unaided eye could resolve details of the beach topography (and therefore also presumably resolve the stark white chests of any pufflings present) with ambient light alone. Observers took turns (of no set duration) using the night vision binoculars to avoid fatigue. The remaining team members scanned the beach, water, and airspace with the unaided eye and recorded each puffling observation, including notes on whether a puffling was suspected or known to be the same individual as one observed earlier. Team members stood next to one another and visually tracked each individual puffling as long as it was within view, continuously communicating about positions and movements of birds to keep track of unique individuals. Whenever visual contact with a puffling was lost, subsequent visual contact with a puffling in the same general location was judged in its likelihood of being the same individual based on the amount of time that had elapsed since previous visual contact. If less than ∼10 seconds elapsed between initial and subsequent visual contact, the second observation was considered to be the same bird; otherwise, it was recorded as a “potential duplicate”.

Any pufflings that became stranded were captured by hand as they were observed and placed individually into a vented plastic holding container (40 cm long × 30 cm wide × 25 cm tall) in a quiet, dark and sheltered location. We took care to limit our time on the beach during capture to a minimum (less than 5 minutes), and only executed a capture attempt when no other birds were seen nearby (either in the water or on land), so as not to influence their behaviour. During a capture attempt, the remaining observer(s) divided their attention amongst scanning the beach, water, and airspace with both night vision binoculars and the unaided eye to ensure all new puffling observations were documented. At the end of the evening (maximum time in captivity, three hours), pufflings were measured and banded by Environment and Climate Change Canada personnel, and released at the water’s edge of the same site where they became stranded (always in dark conditions).

#### Data extraction

From the recorded observational data, we extracted minimum, maximum, and best-estimate counts of unique individual pufflings observed each night at each beach. *Minimum counts* for each beach, on each night were calculated by summing the number of observations known to be of unique individuals, omitting all observations that were potentially duplicates; this was our most conservative count. *Maximum counts* were calculated by summing the number of observations known to be of unique individuals plus all those that were potentially duplicates. *Best-estimate counts* were calculated by summing the number of observations known to be of unique individuals plus only the potential duplicates that were (according to qualitative field notes) likely to be unique individuals; this count is likely the most accurate. On six of eleven nights, the number of pufflings present at the illuminated beach at any one time was low enough to facilitate continuous visual tracking of all or nearly all individuals, resulting in few (if any) potential duplicates and minimum, maximum, and best-estimate counts that vary little from one another. On the other five nights, numbers were high enough that continuous visual tracking of all individuals was not possible despite our best efforts; this is when the number of potential duplicates was highest, and the greater difference between minimum and maximum counts on these nights reflects this difficulty. We also tabulated on a per-treatment basis the number of pufflings observed in the water, in the air, and on the beach; however, it is important to note that because of the difficulty in tracking unique individuals (see above), these are counts of observations rather than estimated counts of unique individuals.

### Response of Pufflings to Light at Fine Spatial Scales

#### Animal collection, housing, and release

To measure phototactic behaviour and activity levels we mainly used stranded pufflings provided by the Puffin Patrol between 8-24 August 2021. Although it is possible that stranded pufflings do not represent a truly random sample of the entire fledgling population, it is this population in which we are most interested since it appears to be most affected by ALAN and also the population from which we would expect the greatest response to our experimental light stimuli. Puffin Patrol volunteers captured pufflings by hand using long-handled butterfly nets, placed them individually into vented plastic holding containers (described above) and then transported them to Puffin Patrol headquarters by vehicle within 2 hours of capture, but usually sooner. CPAWS-NL staff recorded the location and time of capture (if the information was not already submitted electronically by the volunteer(s)) and placed the crated pufflings in a quiet, dark and sheltered location.

Each night between ∼2000 and 2100 hours, we brought the first rescued pufflings in their crates to a grassy area next to our “field labs,” approximately 25 metres away. Pufflings rested in their crates for a minimum of 10 minutes before being brought into the lab and participating in the choice experiment described below. Pufflings were then returned to their crates and allowed to rest outside at ambient temperature for a minimum of 10 minutes before beginning the open field test in the second field lab. Once the open field test was completed, pufflings were returned to the Puffin Patrol’s designated holding area. All pufflings were then measured, banded, and released the following morning by Environment and Climate Change Canada personnel.

In 2022, we performed one additional choice experiment combination (we did not do an open field test in 2022). We captured stranded pufflings between 10-14 August 2022 in the same focal search area as that of the Puffin Patrol, placed individuals in crates, and brought them to Saint Michaels by vehicle (20 minutes travel). Pufflings rested in their crates in a small, dark, quiet room (∼20°C) for a minimum of 10 minutes before we placed them in the choice experiment. After the experiment, we returned pufflings to their crates and released them the following morning from a nearby beach.

#### Materials for choice experiment to measure phototactic behaviour

We used a Y-maze to measure phototactic behaviour and assess pufflings’ preferences for various light spectra (Figure 2; Appendix Figure A2). The metal “acclimation box” (height: 15 cm) opened into the “main box” (height: 22 cm), a large, opaque, blue Rubbermaid container, via a vertically sliding door. Attached to the main box were two adjacent and parallel “choice arms” constructed of 15 cm-diameter thick-walled expandable blue dryer vent hose, each of which led to a small, opaque, blue Rubbermaid container (“choice box”; height: 18 cm). In the middle of the terminal end of each choice box we cut a round hole, approximately 3 cm in diameter and with its centre approximately 8 cm above the ground, which allowed light from the light source to enter the Y-maze. Except for this hole, we covered the exterior light-facing side of each choice box with aluminum foil to prevent light from passing through the wall of the choice box and thereby affecting the spectrum of light visible inside the Y-maze. Further, we placed a 100 × 50 cm Masonite sheet between the two choice arms and choice boxes to eliminate crossover of the different light types (except in the Bright HPS vs. Dim HPS group, where the incident light crossing over from the Bright HPS choice box was used for the Dim HPS choice box). Another small hole (1 cm in diameter) was cut into the outer-most sides of both choice boxes to allow an infrared baby monitor camera (VTech, VM5262-2; British Columbia, Canada) to view the doorway between each choice arm and its corresponding choice box, allowing us to detect when an individual had made a choice (Figure 2).

**Figure 2.**
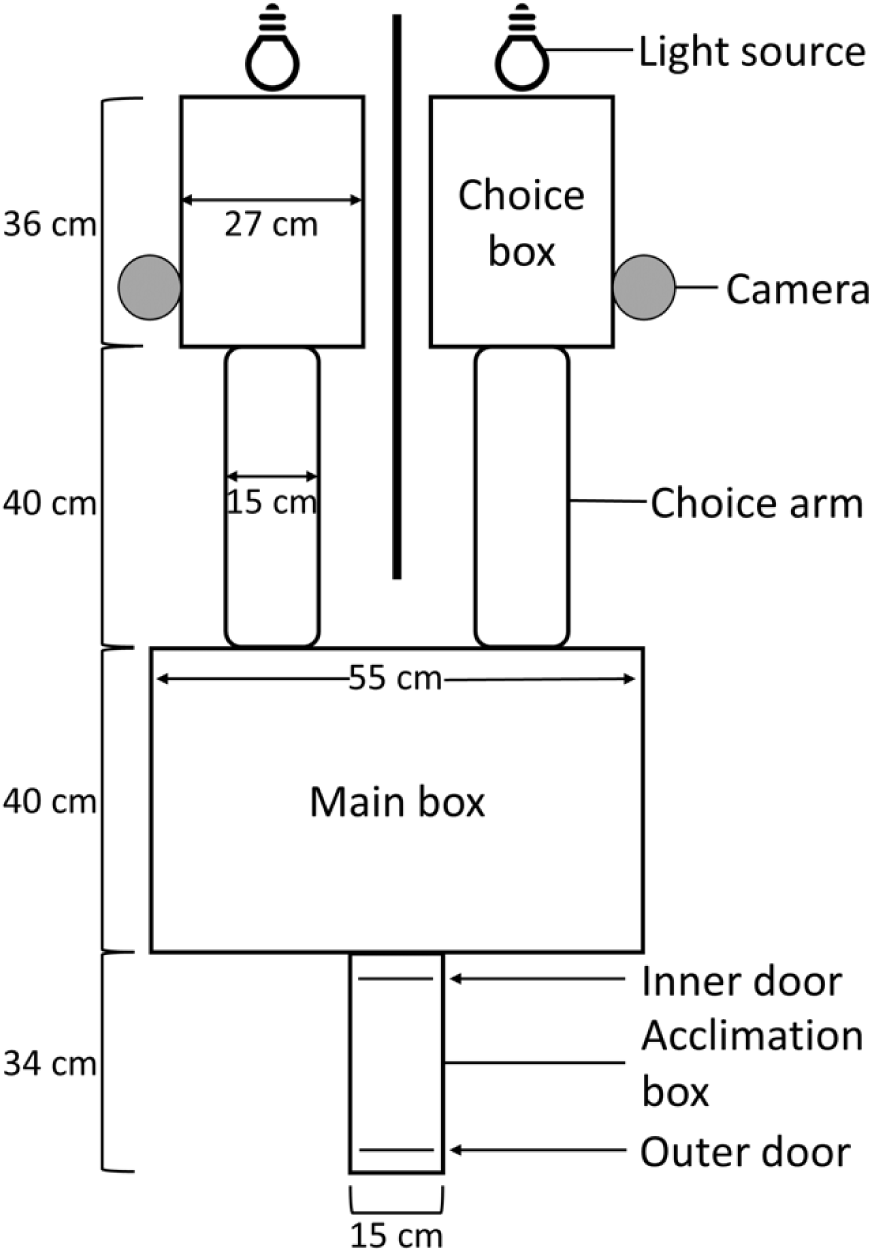
View from above the Y-maze apparatus used for our choice experiment. We placed an individual puffling inside the acclimation box prior to the start of a behavioural trial. The inner door was then opened, and the bird was allowed to enter and explore the main box, choice arms, and choice boxes. An infrared camera detected when an individual became visible at the doorway of a choice box. Note that the diagram is not to scale.

Three different types of lights were used in combination with various filters to produce six spectral choice combinations (spectra displayed in Appendix Figure A2): HPS with a correlated colour temperature (CCT) of 2100 K (Light EnerG, 400 Watt; manufacturer location unknown); Warm White LED with a CCT of 2700 K (Philips, 5W / 40W-equivalent Soft White LED); and Cool White LED with a CCT of 5000 K (Philips, 5W / 40W-equivalent Daylight LED). In total, there were six choice combinations, with number of trials for each bracketed: 1) Bright HPS vs Darkness (*N* = 15); 2) Bright HPS vs Dim HPS (*N* = 22); 3) Bright HPS vs Warm White LED (*N* = 23); 4) Warm White LED vs Cool White LED (*N* = 21); 5) Dim Blue LED vs Orange LED (*N* = 25); 6) Bright Blue LED vs Orange LED (*N* = 25, in 2022 only).

When used, LED lights were fixed to a retort stand and positioned approximately 5 cm from the choice box’s distal hole. The HPS light fixture had an attached rectangular metal reflector on one side, which was bent around the lightbulb to create a cylindrical shape to focus the light out of the distal end of the cylinder. When HPS light was used, the end of the light reflector was placed approximately 15 cm away from the hole. These distances were calibrated with the use of variable neutral density filters (crossed sheets of polyvinyl alcohol-iodine polarizer films; Alight Polarizers, PF006 linear polarizer; Texas, USA) and diffusers (parchment paper) to ensure that the amount of light entering the Y-maze at the peak wavelength of each light type’s respective spectrum was approximately equal across light types, as measured by a calibrated spectrometer (see Appendix Figure A2). The two blue LED light options and the Orange LED light option were created by pairing the Cool White and Warm White LED lightbulbs, respectively, with neutral density filters and diffusers as well as coloured acrylic (ePlastics; California, USA) and glass filters (see Appendix Table A3). We tested pufflings in blue and orange LED light in an attempt to distinguish effects of “warm”- and “cool”-coloured light on phototaxis, following distinctions reported in the literature for other seabird species (Salamolard et al., 2001 as cited in Minatchy, 2004; Rodríguez et al., 2017b).

#### Procedure of choice experiment to measure phototactic behaviour

In 2021, 107 pufflings were randomly chosen from those that were stranded and rescued by the Puffin Patrol and divided among five choice combinations. We tested only one combination each night via selection out of a hat (i.e., it took five consecutive nights to conduct trials in all five combinations, after which they were drawn nightly again in random order), except in the last few days of the field season when we selected the choice combination each night to correct for sample-size deficiencies (all choice combinations tested over a minimum of two non-sequential nights). In 2022, 25 pufflings were tested in Bright Blue LED vs Orange LED, resulting in a total choice experiment sample size of *N* = 132.

Before each trial, the light stimuli were put in their positions and turned on. Each puffling was brought into the field lab, removed from its holding container, and placed in the acclimation box of the Y-maze. After precisely 2 minutes the inner sliding door was removed and the puffling was allowed 13 minutes to make a “choice”. If the puffling did not voluntarily exit the acclimation box by 5 minutes after removal of the inner door, it was gently prodded with the experimenter’s hand reaching through the outer sliding door until the puffling entered the main box. In total, 20% (27 of 132) of pufflings needed prodding. A choice was considered to have been made when a bird reached the end of a choice arm and any part of the body was visible on the monitor (even if it did not fully enter the choice box). The bird was removed from the apparatus after making a choice or after 15 minutes had elapsed (deemed “No Choice”), whichever came first, and returned to its holding container in preparation for the open field test. Between each individual’s trial, the entirety of the Y-maze was cleaned with 70% isopropyl alcohol and a disposable cloth to eliminate possible olfactory cues, and the two lights were alternated between the two choice boxes to account for any potential side bias on part of the birds.

#### Materials to measure activity levels under various light spectra

The cube-shaped open field test arena was constructed from 6 semi-opaque white twin wall polycarbonate panels (1 m × 1 m), held together by acrylonitrile butadiene styrene (ABS) strips. The floor was left unattached for easy removal and cleaning between trials. A 20 cm × 20 cm grid composed of 25 squares (i.e., 5 squares × 5 squares) was created on the floor using black electrical tape. A flap door (approximately 20 cm wide × 15 cm tall) was cut into the bottom edge of one side of the arena to allow pufflings to be placed into and removed from the apparatus. To eliminate any ambient light, an opaque blackout curtain was placed over the roof and sides of the arena. Two holes were cut in the center of the roof and overlaying curtain: one hole, approximately 3 cm in diameter, accommodated the lens of a miniature action camera (Activeon, CCA10W, Activeon Inc.; San Diego, CA, USA) with a wide-angle lens to allow for full view of the floor of the arena. The other hole, approximately the same size and 5 cm away, allowed for a controlled amount of light to enter the arena from an overhanging light source. When in use, LED lights rested directly on the hole, while the HPS light was suspended approximately 20 cm above the hole. This was done to decrease the amount of heat applied to the arena roof (since the HPS light was warmer than the LED lights), and because the HPS light was brighter than the LED lights (this distance resulted in a relatively equal amount of light entering the arena). Median illuminance was similar across the three light treatments when measured at 25 different locations at floor level (Kruskal-Wallis test: *H*_2_ = 1.3696, *P* = 0.5) using a spectrometer (SRI-2000 Spectral Light Meter; Allied Scientific Pro; Gatineau, Quebec, Canada). The median illuminance from HPS was 26.35 lux (IQR 21.08 – 33.18), while that from Warm White LED was 24.34 lux (IQR 20.30 – 27.80), and that from Cool White LED was 23.88 lux (IQR 19.46 – 28.62). These median illuminance values are comparable to that of a similar behavioural study, wherein illuminance was approximately 30 lux (Atchoi et al., 2023).

#### Experimental procedure to measure activity levels under various light spectra

To determine whether puffling activity level was affected by light type, we performed an open field test. In 2021, 96 of the 107 Atlantic Puffin fledglings used in the choice experiment were used in the open field test, whereby we exposed them to either darkness or one of three light spectra. Five individuals were removed from the final dataset due to camera malfunction, resulting in a final sample size of *N* = 91. Treatments were: 1) Darkness (*N* = 23); 2) HPS (*N* = 19); 3) Warm White (2700 K) LED (*N* = 24); and 4) Cool White (5000 K) LED (*N* = 25). We did not measure activity in response to blue or orange light to focus our testing on spectra from light sources currently used in human settlements (i.e., HPS and LED). Each night, we used only two light treatments in the open field test, and they were never the same as the two light types used in the Y-maze on the same night. The initial treatment each night was chosen randomly, and treatments were alternated for the rest of the night.

Prior to each open field trial, the appropriate light was set up and turned on (except in the Dark treatment), and a video recording (with audio) was started. The puffling was placed just inside the flap door of the arena, and an electronic timer started. After 10 minutes, the video and audio recording were stopped. The puffling was removed from the arena, replaced in its holding container, and returned to Puffin Patrol headquarters. The floor of the arena was not cleaned between trials unless a puffling defecated during its trial, in which case the affected area was cleaned with 70% isopropyl alcohol and allowed to dry before the next trial. As many trials as possible were completed each night (median 10 trials; range 3-13 trials per night).

#### Activity level data extraction

Prior to reviewing any open field test videos, we determined the order in which to watch each video using the random number “Sequence Generator” on random.org (Randomness and Integrity Services Ltd., Dublin, Ireland). We used behavioural scoring software JWatcher (Version 1.0, Macquarie University and University of California, Los Angeles) to score two levels of behaviour for 10 minutes: “mobile” behaviour included walking, running, or flying; otherwise, birds were considered “immobile”. The software automatically calculated the total amount of time spent mobile and immobile by each puffling in milliseconds, based on the timing of our key presses, and we subsequently converted each score of time spent mobile into seconds.

In the case of “Dark” treatment videos, only the audio was discernible. Puffins have large, webbed feet and claws, which allowed us to hear them as they walked over the polycarbonate floor of the arena. Instead of viewing the file, we listened to its audio and recorded instances when we heard the bird moving (mobile) or halting (immobile). Billing (tapping or scraping the arena walls with the bill) and scratching (clawing at the arena floor with the feet) could also be heard but were not scored as “mobile” behaviour on their own. Each time a key code was pressed, the previous behaviour was automatically assumed to have ceased. We decided to score only “Dark” videos audially (rather than scoring all videos audially) because: 1) we reasoned that visual scoring is both more accurate (i.e., truer to the correct value) and more precise (i.e., repeatable) than audial scoring, and is therefore preferable when three of our four treatments can be scored visually; 2) the noise associated with the HPS light’s ballast fan would have made audial scoring of those videos difficult and therefore introduce a new source of scoring bias; and 3) extra care was taken during the Dark treatment specifically to ensure that noises similar in quality to those of puffling footsteps were reduced to an absolute minimum so as to increase the accuracy of those audial scores, whereas only reasonable care was taken to reduce similar noises in the lighted treatments.

To determine whether the time spent mobile and immobile in the Dark treatment could be accurately scored by listening to the audio alone, we selected a subset of 25 videos from the three lighted conditions (HPS, Warm White LED, Cool White LED) that had already been scored. We listened to all 25 videos in random order with headphones and scored them audially as normal. A single observer did all visual and audial scoring.

#### Statistical Analysis

All analysis was performed in program R (ver. 4.3.0; R Core Team, 2021). We report results to a significance level of *α* = 0.05.

#### Response of pufflings to light at a coarse spatial scale

We tabulated the *minimum*, *maximum*, and *best-estimate counts* of pufflings seen during the “Light” and “Dark” treatments each night, as well as counts of observations of pufflings in the water, in the air, and on the beach. To control for search effort, we calculated “estimated observation rate” (reported as birds per observer) by dividing the best-estimate counts of pufflings during a given treatment on a given night (over the two-hour experimental period) by the number of observers present. We report medians and interquartile ranges. For a more conservative approach we also compared median estimated observation rates on the beach only (estimated count on the beach divided by number of observers), between Light and Dark. This was because the night vision binoculars worked best while looking at a solid backdrop (i.e., the beach itself) and because our land-based search area at both beaches was standardized. The results of this experiment were definitive in demonstrating higher numbers of pufflings at the beaches during the “Light” treatment regardless of the metric used, obviating the need for formal statistical analysis.

### Response of pufflings to light at fine spatial scales

#### Phototactic behaviour

In the choice experiment we tested a final sample size of *N* = 131 pufflings. To test whether pufflings displayed evidence for positive phototaxis or preferences for certain light spectra over others we excluded individuals who exhibited No Choice (*N* = 23) since this behavioural outcome indicates no preference. For the remaining individuals, we performed a binomial test (package “*stats*”, v. 4.3.0) of the light stimuli chosen in each of the six choice combinations to determine if choices significantly differed from hypothesized proportions of 0.5, and we report 95% confidence intervals of the estimated true proportions. We then conducted a Fisher exact test (package “*stats*”, v. 4.3.0) on the number of pufflings from each combination that were “prodded” to determine if the proportion of prodded individuals differed among combinations.

#### Activity levels

To determine if scoring videos audially may have artificially biased our estimates of time spent mobile in the “Dark” treatment group (in comparison to scoring the light videos, which was done visually), we conducted a simple linear regression of time spent mobile scored visually as a function of time spent mobile scored audially of the 25 selected lighted videos that were scored both visually and audially. The regression equation was statistically significant (*r*^2^ = 0.985, *F*_1,23_ = 1478, *P* < 0.001); audial scores slightly overestimated but nonetheless strongly predicted visual scores of time spent mobile (β = 0.9228, *P* < 0.001, Appendix Figure A3). We converted the “Dark” data using the regression equation and used these “converted” values in statistical comparisons with the unmodified visual scores from the other treatment groups (see Appendix). Note that the average difference between raw and converted scores of time spent mobile in the “Dark” group was a decrease of only eight seconds (maximum difference, −21 seconds).

We identified one outlier in the entire dataset: an individual exposed to Warm White LED light that spent 353 seconds mobile (a deviation from that treatment group’s 75% quartile by approximately +2.3 times the interquartile range). Because it is unclear what caused this individual to behave so differently from the rest of the individuals exposed to Warm White LED light, we removed it from all analyses (both choice experiment and open field test). Retention of this outlier causes a loss of statistical significance in one out of three comparisons, but the general trends remain unchanged (to see results with the outlier included, see Appendix). Because activity level data did not meet assumptions of normality or homogeneity of variance, we used non-parametric statistics. To test whether individuals differed in the time spent mobile across treatment groups, we performed a Kruskal-Wallis H test (package “*stats*”, v. 4.3.0) and a post hoc Dunn test (package “*FSA*”, v. 0.9.4; Ogle et al., 2023) for multiple comparisons.

## RESULTS

### Response of Pufflings to Light at a Coarse Spatial Scale

We calculated a best-estimate total of 136 pufflings that arrived at the beaches (observed in the water, on the beach, and in the air combined) during the Light treatment, and only 2 pufflings during the Dark treatment, over the 11 nights of the experiment. Similar patterns emerged between the Light and Dark treatments when we expressed best-estimate count data on a nightly basis (Figure 3A): during the Light treatment the median best-estimate count was 7.0 (IQR 4.5 – 14.5) individuals per night compared to 0.0 (IQR 0.0 – 0.0) in the Dark. When numbers are compared within the same experimental night, *minimum*, *best estimate*, and *maximum* counts of pufflings observed, as well as observation rates that control for search effort, all tell a similar story: the frequency of pufflings was consistently higher during Light treatments than during Dark treatments (Appendix Tables A1, A2). When we controlled for search effort, median estimated observation rate was 2.0 (IQR 1.2 – 6.0) birds per observer in the Light and 0.0 (IQR 0.0 – 0.0) birds per observer in the Dark (Figure 3B). Focussing only on the standardized 50-metre-long beach search areas for which we have greatest confidence, pufflings were observed 54 times during the Light treatment, which greatly outnumbered the Dark treatment’s single observation. Controlling for search effort, a median of 0.7 birds per observer (IQR 0.4 – 2.3) was seen on the beach in the Light treatment, compared to 0.0 birds per observer (IQR 0.0 – 0.0) in the Dark. Behavioural observations indicated that pufflings in flight generally flew at altitudes of 15 metres or less, in straight lines either along the shoreline (in both directions) or overhead going inland, and pufflings in the water and on the beach approached the light to varying distances (approximately 15-40 metres) and then either stopped moving (to preen, rest, etc.) or proceeded back to the water and swam away (if not captured first).

**Figure 3.**
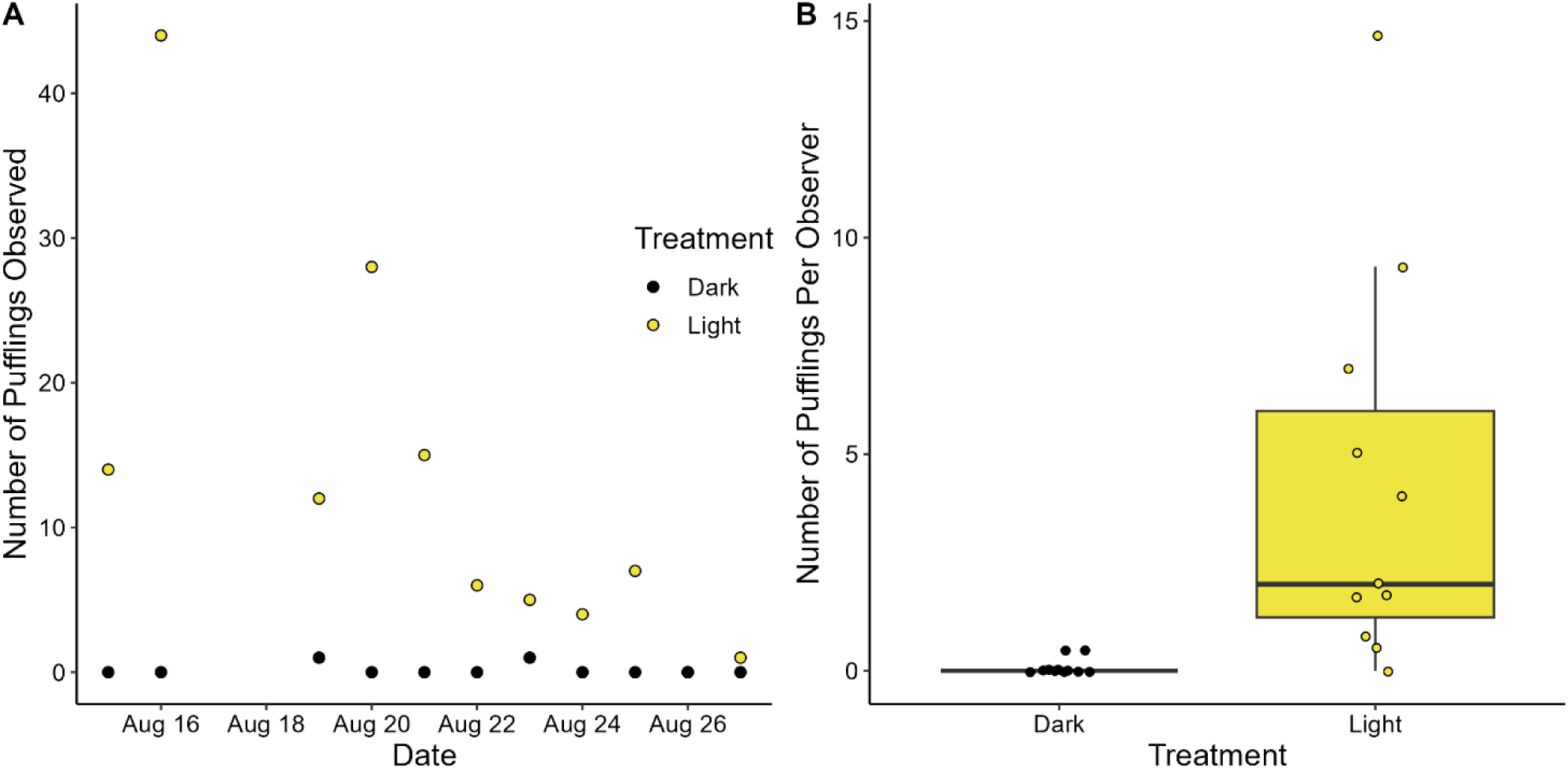
Number of fledgling puffins observed per night at artificially lit and dark beaches. A) Best-estimate total counts of pufflings (in the water, on the beach, and in the air combined) seen each night during the Dark (black circle) and Light (yellow circle) treatments, 15-27 August 2022. Note that points overlap on August 26 due to zero birds being observed during either treatment. B) Observation rates (same best-estimate counts of pufflings observed, divided by the number of observers present) at the two beach sites during Dark and Light treatments. Each data point represents the observation rate on one experimental night at each site and points have been offset from one another to improve readability.

### Assessment of Phototactic Behaviour and Light Preferences

Significantly more individuals chose Bright HPS light than Dark (*P* = 0.002; 95% CI of estimated true proportion of Bright HPS: 0.69-1.00; Table 1) in the choice experiment. Although not statistically significant, more individuals chose Bright HPS over Dim HPS (*P* = 0.1; 95% CI of estimated true proportion of Bright HPS: 0.46-0.88). There was no significant preference of any light type in any other remaining choice combination (Table 1). There was no statistical evidence for side bias, as pufflings chose Choice Box 1 (*N* = 57) relatively equally to Choice Box 2 (*N* = 52; binomial test: *P* = 0.7; 95% CI of estimated true proportion of Choice Box 1: 0.43-0.62). There was no difference among choice combinations in the number of pufflings prodded out of the acclimation box after 5 minutes (Fisher Exact Test: *P* = 0.6).

**Table 1.** Number of pufflings that chose each of the two light options in each choice combination. Individual binomial tests assessed evidence for a preference of one light type over the other in each choice combination (excluding “No choice” individuals), using 0.5 as the theoretical (null) proportion for each of the two options. We report the 95% confidence intervals of the true proportion of “Light type 1” as a reference. We also present the number of individuals that made “No choice” in each combination.

| Light type 1 | Quantity | Light type 2 | Quantity | 95% CI | *P*-value | No choice |
| --- | --- | --- | --- | --- | --- | --- |
| Bright HPS | 10 | Dark | 0 | 0.69-1.00 | 0.002 | 5 |
| Bright HPS | 14 | Dim HPS | 6 | 0.46-0.88 | 0.1 | 2 |
| Bright HPS | 10 | Warm White | 11 | 0.26-0.70 | 1.0 | 2 |
| Warm White | 10 | Cool White | 10 | 0.27-0.73 | 1.0 | 2 |
| Dim Blue | 13 | Orange | 9 | 0.36-0.79 | 0.5 | 2 |
| Bright Blue | 9 | Orange | 6 | 0.32-0.84 | 0.6 | 10 |

### Assessment of Activity Levels in Different Light Types

In our open field test, there was a significant effect of light type on time spent mobile by pufflings (Kruskal-Wallis test: *H*_3_ = 18.396, *P* = 0.0004). Pufflings spent more time mobile in Darkness (median time spent mobile = 120 seconds) and in HPS light (median 111 seconds) than in Cool White LED light (median 32 seconds; Dunn test: *P* = 0.003 and *P* = 0.004, respectively; Figure 4). Pufflings also spent more time mobile in Darkness than in Warm White LED light (median 54 seconds; Dunn test: *P* = 0.045). Pufflings spent more time mobile, but only marginally, in HPS light compared to Warm White LED light (Dunn test: *P* = 0.050; Figure 4). If the statistical outlier is included (see Methods), the difference between Warm White LED and Darkness is no longer significant, although the trend remains the same (*P* = 0.09; see Appendix).

**Figure 4.**
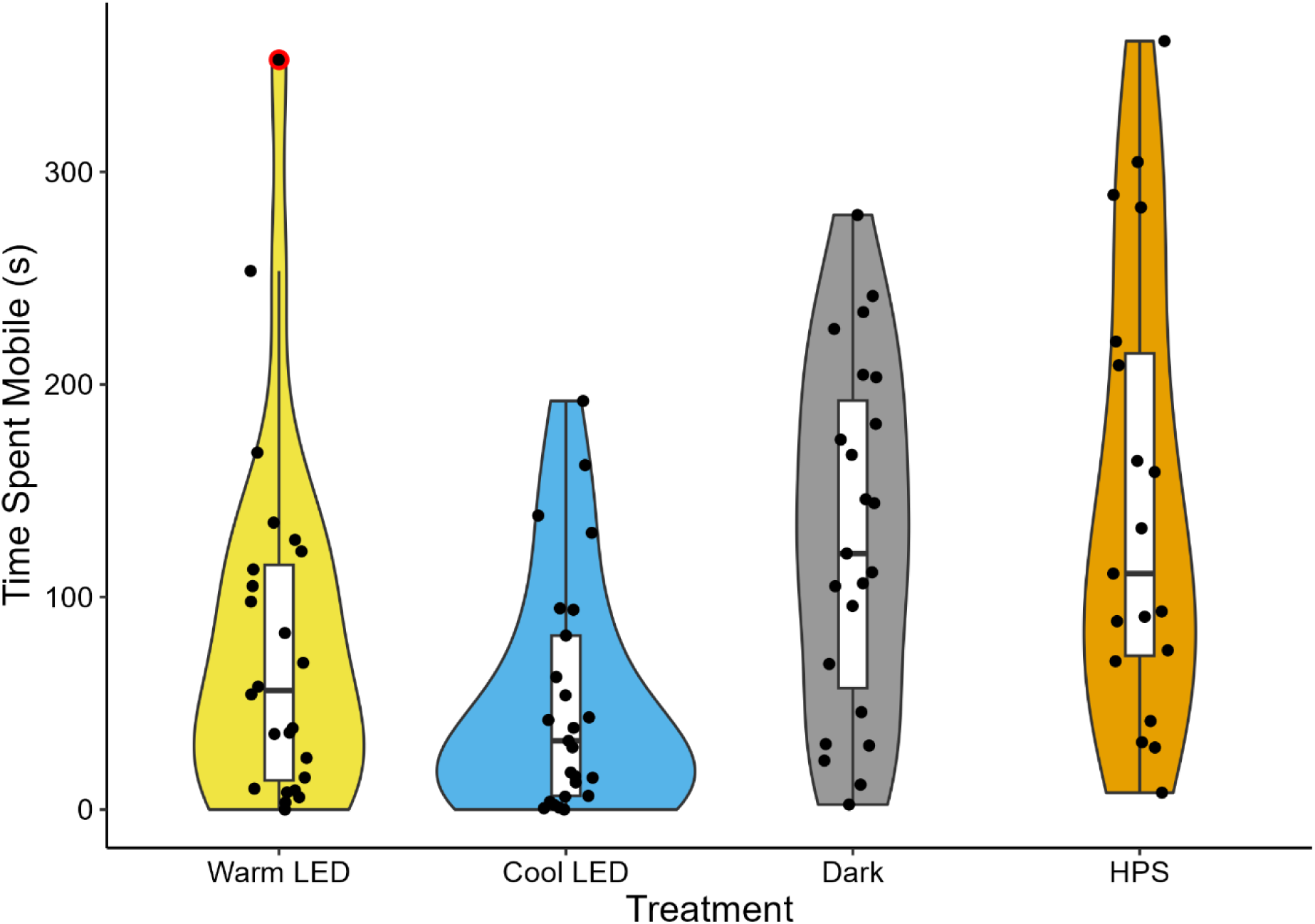
Time spent mobile by fledgling puffins when exposed to each of four light treatment groups. Figure combines a violin plot and boxplot with overlaid individual data points. Sample sizes: Warm White LED light (*N* = 24), Cool White LED light (*N* = 25), Darkness (“converted” data; *N* = 23, see text), and High Pressure Sodium (HPS) light (*N* = 19). The single data point >300 seconds in the warm LED treatment (outlined in red) was considered a statistical outlier and was not included in analysis. Individuals were significantly less mobile when exposed to Warm White LED and Cool White LED than when exposed to Darkness, and significantly less mobile under Cool White LED than under HPS.

## DISCUSSION

Our results form the first experimental evidence that light attraction behaviour is a causal mechanism behind the stranding of Atlantic Puffin fledglings (pufflings) in coastal areas during their initial fledging flights. This evidence emerges at both coarse and fine spatial scales. Even after controlling for search effort and size of the search area, in our coarse-scale beach illumination experiment we found significantly more pufflings stranded during the Light treatment than during the Dark treatment. Further, consistent with a light attraction hypothesis of stranding, in our y-maze experiments pufflings significantly preferred light over darkness and somewhat preferred bright HPS light over dim HPS light (although the latter was not statistically significant). Contrary to our expectation that pufflings would prefer “cool” light spectra over “warm” light spectra (based on results of Salamolard et al., 2001 as cited in Minatchy, 2004; Rodríguez et al., 2017b), we found no preferences when pufflings were simultaneously presented with two different light spectra. Our open field test experiment, in which we exposed pufflings to either darkness or one of three light types, revealed that pufflings were more active in Darkness than in any LED light, and more active in HPS light than under Cool White LED light. Pufflings also tended to be more active in HPS light than in Warm White LED light, but this difference only bordered on statistical significance.

We are first to demonstrate a reduction in stranding numbers of an alcid through the experimental reduction of ALAN. Our finding that more pufflings became stranded at our two beach sites during the Light treatment compared to the Dark treatment is consistent with similar studies on procellariiforms (Reed et al., 1985; Miles et al., 2010; Rodríguez et al., 2014; Burt et al., 2024). Our results therefore provide further support for recommendations to reduce or eliminate sources of ALAN near breeding colonies of seabirds affected by stranding, especially where populations are already in decline (e.g., Rodríguez et al., 2017a). However, it should be noted that the number of pufflings stranded during our experiment, and indeed during the entire Puffin Patrol rescue program, is small relative to the estimated total fledging cohort (e.g., in 2019 an estimated 246,069 ± 25,988 juveniles successfully fledged from the Witless Bay Ecological Reserve and 326 were found stranded; Wilhelm et al., 2021). It is possible, then, that navigation error during natural dispersal movements inevitably brings a small proportion of fledglings close to land, where they become attracted to and stranded by nearby artificial lights (Brown et al., 2023).

Previously, we expressed concern that human visual perception is more acute, and therefore the detection of stranded birds is likely more efficient in light conditions compared to darkness, potentially leading to a bias in detection probability (Brown et al., 2023). As such, in our coarse-scale light attraction experiment we attempted to reduce the bias between our two treatments by: 1) using identical night vision binoculars at each of our sites on each night of the experiment; 2) clearly delineating a search area of the same size (area of approximately 50 m × 15-20 m) on both beaches that took into account the maximum visual range of the binoculars; and 3) controlling for the number of observers present at each site in our data analysis. Even if our Light treatment was still subject to bias toward elevated detection probability, it is highly unlikely that the large difference in number of pufflings found within the beach search areas (*N* = 54 in Light vs *N* = 1 in Dark) is attributable to missing a significant number of pufflings in the Dark treatment. In fact, the single puffling we detected within the search area during the Dark treatment was on a night with very low lunar illumination (13.3%), after the moon had already set, and during overcast and rainy conditions. That pufflings were visible to observers even in the worst of visibility conditions suggests that if more individuals were indeed present in the Dark treatment, they would have been detected. Additionally, by alternating each night the site location for Light treatment, we simultaneously controlled for the potential effects of date, moon phase (see Wilhelm et al., 2021) and site on the numbers of pufflings we observed.

In addition to the stark difference in number of pufflings observed within the delineated beach search areas in the Light versus Dark treatment, more individuals were also observed swimming and flying in the Light treatment (*N* = 77 and *N* = 78, respectively) than in the Dark treatment (*N* = 1 and *N* = 1, respectively).

Support for the light attraction hypothesis of stranding in pufflings is bolstered by the positive phototactic behaviour pufflings displayed in our choice experiment. Pufflings chose Bright HPS light significantly more often than darkness and chose Bright HPS light slightly (but insignificantly) more often than Dim HPS. These results agree with those of a similar study in which Ancient Murrelet chicks chose light significantly more often than darkness in a T-maze just after leaving their nests (Gaston et al., 1988), but interestingly are contrary to results of a Y-maze experiment in which Cory’s shearwater fledglings chose darkness over light (Atchoi et al., 2024). Ancient Murrelet chicks leave the nest at night under parental care on average only two days after hatching (Gaston and Shoji, 2020), long before they are capable of flight. It is this lack of flight ability, combined with breeding in mostly remote areas away from human settlement, that presumably precludes them from stranding in areas with ALAN. That our Atlantic puffin Y-maze results are consistent with those of Ancient murrelets but not Cory’s shearwaters possibly suggests a taxonomic difference in fine-scale behavioural responses to ALAN between the two main seabird groups affected by light-induced stranding (i.e., alcids and procellariiforms); however, more research is necessary to confirm this trend.

Contrary to our expectations, pufflings did not display any preferences for certain light spectra over others when presented with a choice. We based our expectations on two premises: first, previous studies have found that attraction behaviour and stranding numbers of fledgling procellariiforms increase under exposure to short wavelength-dominant (i.e., high CCT) light types like metal halide (∼4500 K CCT) compared to long wavelength-dominant (low CCT) light types like high pressure sodium (∼2000 K CCT; Salamolard et al., 2001 as cited in Minatchy, 2004; Rodríguez et al., 2017b), although this is not always the case (e.g., shielding of long wavelength light from radiating upward may negate the effect; Urmston et al., 2022; Atchoi et al., 2024) and evidence is limited. Second, the procellariiform eye is quite sensitive to short wavelengths (Reed, 1986; Bowmaker et al., 1997; Hart, 2004), implying a potential link between the wavelengths of highest spectral sensitivity and those that induce the strongest phototactic responses. Atlantic Puffins likely have similar spectral sensitivities given the similarities in photoreceptor profiles between auks and procellariiforms (Ödeen and Håstad, 2003), which is why we expected they would choose blue light over orange and cool white over warm white LED light in our Y-maze. It is therefore unclear why these choice combinations did not elicit differences in phototactic behaviour. Most of our light stimuli were calibrated such that the amount of light irradiated at the peak wavelength of each spectrum was relatively equal; we did this because the calibration of spectra based on perceived “brightness”, although valuable, is difficult without a complete understanding of spectral sensitivities in the species of interest. To a puffling’s eyes our provided light options were therefore not necessarily perceived as all being of the same “brightness”, and there may thus have been unquantified effects of perceived brightness on the choice behaviour they exhibited in some of our choice combinations. Another possible explanation is that the locomotory context of the birds (i.e., standing and walking) and/or the spatial scale of the experimental apparatus are so contextually different from flying in a three-dimensional airspace that they elicit a completely different mode of behaviour as it relates to lighting conditions. A coarse-scale beach illumination experiment that compares phototactic responses and stranding numbers across several different light spectra (ideally calibrated according to perceived brightness) may provide insight in a more realistic context (as per Rodríguez et al., 2017b).

When we measured the activity levels of pufflings exposed to darkness and various spectra of artificial light in our open field test, they spent significantly more time active in Darkness than in Cool or Warm White LED light (despite the effect of “converting” the Dark treatment scores to reduce the significance of these differences), and significantly more time active in HPS light than in Cool White LED light. It is unclear what factors drove our results, especially in Darkness. It is possible that in the HPS light treatment the low-level noise of the associated ballast fan (unlike the LED lights which have no ballast fan) elicited an attractive response, which has been anecdotally observed in stranded pufflings (Harris, 1982; Harris et al., 1998; reviewed in Wilhelm et al., 2013). Conversely, it is possible that the greater proportion of cool-hued light in LED spectra compared to HPS has a calming effect on pufflings that results in reduced activity, as blue light does in domestic ducks (Sultana et al., 2013) and in Black-headed Buntings (*Emberiza melanocephala*) experiencing periods of migratory restlessness (but with this pattern reversed outside these periods; Yadav et al., 2015). A fear-based freezing response toward some aspect of LED light could also drive the reduced activity levels we observed in that light type, similar to that observed in some poultry lines when placed in an open field arena (Forkman et al., 2007; Campbell et al., 2019), although our anecdotal observations of pufflings preening themselves in the arena during bouts of immobility would appear to contradict this hypothesis. Ultimately, the true cause of the reduced activity levels that we observed in pufflings exposed to LED light, especially in Cool White LED light, remains a matter of conjecture. Regardless of the reason, these differences in activity patterns may hold implications for detection probability following stranding; for example, pufflings may be easier to find in darkness and under HPS light if they are more mobile there, whereas comparatively less active birds under LED light may be less noticeable, all else being equal.

### Conservation Implications

Globally, there is a concerted effort to reduce the impact of ALAN on wildlife, which includes efforts to mitigate and prevent the stranding of various seabird species. To date, most research has focused on procellariiforms, with comparatively little attention paid to the Atlantic Puffin. We have demonstrated experimentally that the addition of sources of ALAN near Atlantic Puffin breeding colonies has the potential to drastically increase the number of fledglings that become stranded, which in turn would almost certainly contribute to decreased juvenile survival. However, perhaps counter-intuitively, we suggest that light could also be used to lure wayward puffins away from areas of high mortality risk (i.e., towns and roads) to those of low risk (e.g., supervised remote sites) as part of a dynamic rescue effort, and this potential application of ALAN in a conservation context deserves further research. From fine-scale behavioural experiments, we demonstrated that Atlantic Puffin fledglings display positive phototactic behaviour in response to all tested bulb colours, without a preference for either blue- or red-dominant spectra. Although there may also be compounding effects of perceived brightness, this implies that changes to street lighting from historically prevalent HPS lamps to more energy-efficient LED, or (if specifically attempting to reduce attraction) from cool-hued to warm-hued lamps, may not be effective in reducing strandings in Atlantic Puffins and that the reduction and shielding of ALAN sources remain the best recommendations. Our open field test results imply that post-stranding LED light exposure may have either a calming or startling effect on stranded pufflings, with potential implications for rescue, rehabilitation, and even husbandry of puffins and other auks.

## Data availability statement

Data are available as Supplementary Material.

## Declaration of Generative AI and AI-assisted technologies in the writing process

During the preparation of this work the authors used ChatGPT to formulate a first draft of the title. After using this tool, the authors reviewed and edited the title as needed and take full responsibility for its content.

## APPENDIX

### Methods

#### Conversion of “Dark” treatment activity levels

To determine if we should “convert” the audial scores of time spent mobile by pufflings in the “Dark” treatment (*x*) to estimated scores of time spent mobile had been scored visually (*y*), we calculated 95% confidence intervals of the slope and intercept estimates (that is, 0.92 and 2.30, respectively). We decided *a priori* that if either the slope estimate’s 95% confidence intervals did *not* contain 1.00 or if the intercept estimate’s 95% confidence intervals did *not* contain 0.00, we would convert the “Dark” data using *y = 2.3 + 0.92x*; otherwise, we would use the raw “Dark” data for analysis. The 95% confidence interval of the slope was [0.87, 0.97] and that of the intercept was [-5.29, 9.95]. The slope estimate’s 95% confidence intervals therefore did not contain 1.00, so we converted the “Dark” data for analysis.

### Results

#### Assessment of activity levels in different light types when outlier is included

The following results were obtained when the individual identified as an outlier (in the Warm White LED light treatment of the open field test; spent 353 seconds mobile) was retained in our analysis. There was a significant effect of light type on time spent mobile by pufflings (Kruskal-Wallis test: *H*_3_ = 17.184, *P* < 0.001); post-hoc testing revealed that pufflings spent more time mobile in Darkness (median time spent mobile = 120 seconds) and in HPS light (median 111 seconds) than in Cool White LED light (median 32 seconds; *P* = 0.004 for both comparisons; Figure 4). Pufflings also tended to spend more time mobile in Darkness and in HPS light than in Warm White LED light (median 56 seconds; *P* = 0.09 and *P* = 0.10, respectively), but these differences failed to reach significance (Figure 4).

### Tables

**Table A1.** Minimum (“Min”), best-estimate (“Est”), and maximum (“Max”) counts of pufflings observed at the two beaches during Dark and Light treatments over the course of 11 experimental nights, accompanied by percent moon illumination on each night (“Moon %”) and qualitative observations of weather conditions at the sites. All counts for both treatments were conducted over the same experimental duration each night (two hours). Percent moon illumination data were collected from moongiant.com. Weather conditions were not recorded at the sites on night 4; conditions on this night were obtained from Environment and Climate Change Canada (climate.weather.gc.ca).

| Night | Moon % | Weather | Dark |  |  | Light |  |  |
| --- | --- | --- | --- | --- | --- | --- | --- | --- |
|  |  |  | Min | Est | Max | Min | Est | Max |
| 1 | 85 | Clear | 0 | 0 | 0 | 10 | 14 | 23 |
| 2 | 76 | Cloud & fog | 0 | 0 | 0 | 34 | 44 | 52 |
| 3 | 47 | Cloud & rain | 1 | 1 | 1 | 11 | 12 | 17 |
| 4 | 38 | Drizzle* | 0 | 0 | 0 | 27 | 28 | 30 |
| 5 | 29 | Clear | 0 | 0 | 0 | 14 | 15 | 19 |
| 6 | 20 | Clear | 0 | 0 | 0 | 6 | 6 | 6 |
| 7 | 13 | Cloud & rain | 1 | 1 | 2 | 5 | 5 | 5 |
| 8 | 8 | Cloud & wind | 0 | 0 | 0 | 4 | 4 | 4 |
| 9 | 3 | Cloud | 0 | 0 | 0 | 6 | 7 | 8 |
| 10 | 1 | Cloud & wind | 0 | 0 | 0 | 0 | 0 | 0 |
| 11 | 0 | Clear | 0 | 0 | 0 | 1 | 1 | 1 |

\*Weather data obtained from Environment and Climate Change Canada for St. John’s International Airport (47.618611°N, 52.752500°W), located 38-40 km from the two beach sites used in this experiment.

**Table A2.** Minimum (“Min”), best-estimate (“Est”), and maximum (“Max”) “observation rates” (pufflings seen, divided by number of observers) at the beaches during Dark and Light treatments over the course of 11 experimental nights. All observation rates for both treatments apply to a two-hour experimental duration.

| Night | Dark |  |  | Light |  |  |
| --- | --- | --- | --- | --- | --- | --- |
|  | Min | Est | Max | Min | Est | Max |
| 1 | 0 | 0 | 0 | 5 | 7 | 11.5 |
| 2 | 0 | 0 | 0 | 11.3 | 14.7 | 17.3 |
| 3 | 0.5 | 0.5 | 0.5 | 3.7 | 4 | 5.7 |
| 4 | 0 | 0 | 0 | 9 | 9.3 | 10 |
| 5 | 0 | 0 | 0 | 4.7 | 5 | 6.3 |
| 6 | 0 | 0 | 0 | 2 | 2 | 2 |
| 7 | 0.5 | 0.5 | 1 | 1.7 | 1.7 | 1.7 |
| 8 | 0 | 0 | 0 | 0.8 | 0.8 | 0.8 |
| 9 | 0 | 0 | 0 | 1.5 | 1.75 | 2 |
| 10 | 0 | 0 | 0 | 0 | 0 | 0 |
| 11 | 0 | 0 | 0 | 0.5 | 0.5 | 0.5 |

**Table A3.** Materials used to create each of the eight light options used in the choice experiment, including the specific lightbulbs used, whether a neutral-density filter or diffuser were used, and any other materials used.

| **Light option** | **Lightbulb** | **NDF<sup>a</sup>** | **Diffuser<sup>b</sup>** | **Other** |
| --- | --- | --- | --- | --- |
| Dim HPS | 2100 K HPS<sup>c,d</sup> | none | yes | none |
| Bright HPS | 2100 K HPS<sup>d</sup> | yes | yes | none |
| Cool white LED | 5000 K LED<sup>e</sup> | yes | yes | none |
| Warm white LED | 2700 K LED<sup>f</sup> | yes | yes | none |
| Bright blue LED | 5000 K LED<sup>e</sup> | none | none | ePlastics #2424 dark blue acrylic; cobalt glass |
| Dim blue LED | 5000 K LED<sup>e</sup> | yes | yes | ePlastics #2424 dark blue acrylic; cobalt glass |
| Orange LED | 2700 K LED<sup>f</sup> | yes | yes | ePlastics #2422 amber orange acrylic |
| Darkness | none | none | none | light hole covered |

<sup>a</sup>A variable neutral density filter (NDF) consisted of two pieces of polyvinyl alcohol-iodine polarizing film, overlapping at varying degrees of polarization depending on the light option.
<sup>b</sup>Diffuser consisted of one piece of parchment paper.
<sup>c</sup>Incident light only (i.e., crossover light from Bright HPS on other choice box).
<sup>d</sup>High pressure sodium; Light EnerG, 400 Watt
<sup>e</sup>Light-emitting diode; Philips, 5W / 40W-equivalent Daylight LED
<sup>f</sup>Light-emitting diode; Philips, 5W / 40W-equivalent Soft White LED

### Figures

**Figure A1.**
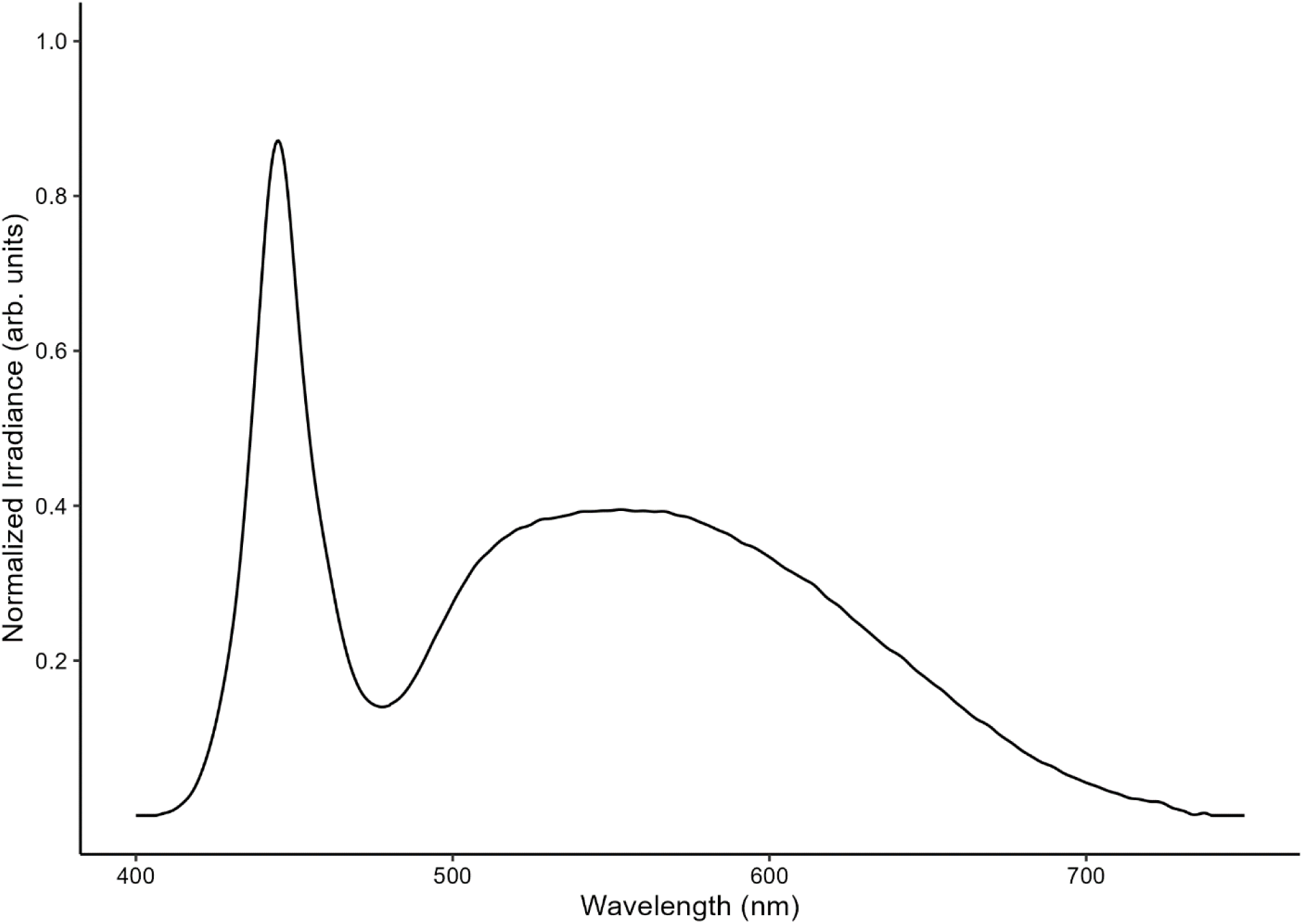
Spectrum of the 6390 K, 30 000 lumen LED light apparatus used on two beach sites to test the responses of pufflings to light at a coarse spatial scale. Note that the y-axis has been normalized to arbitrary units with a maximum of 1.0.

**Figure A2.**
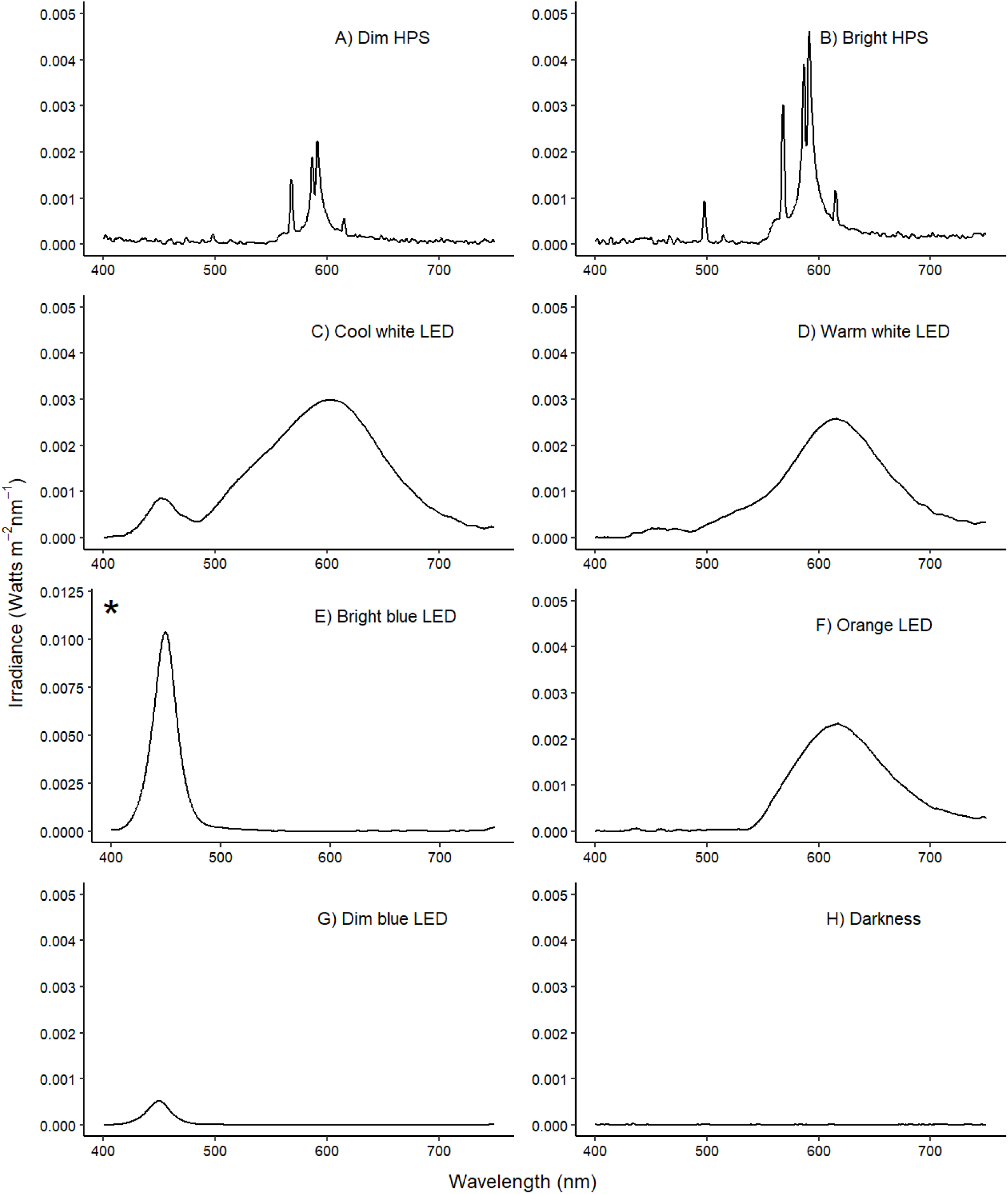
Spectra of all light options used in the choice experiment: A) Dim HPS; B) Bright HPS; C) Cool White (5000 K) LED light; D) Warm White (2700 K) LED light; E) Bright Blue LED light; F) Orange LED light; G) Dim Blue LED light; and H) Darkness. Irradiance is as measured by a spectrometer with its sensor placed at the entrance of the “choice arm”, inside the “main box” of the Y-maze. * = note the change in y-axis scale.

**Figure A3.**
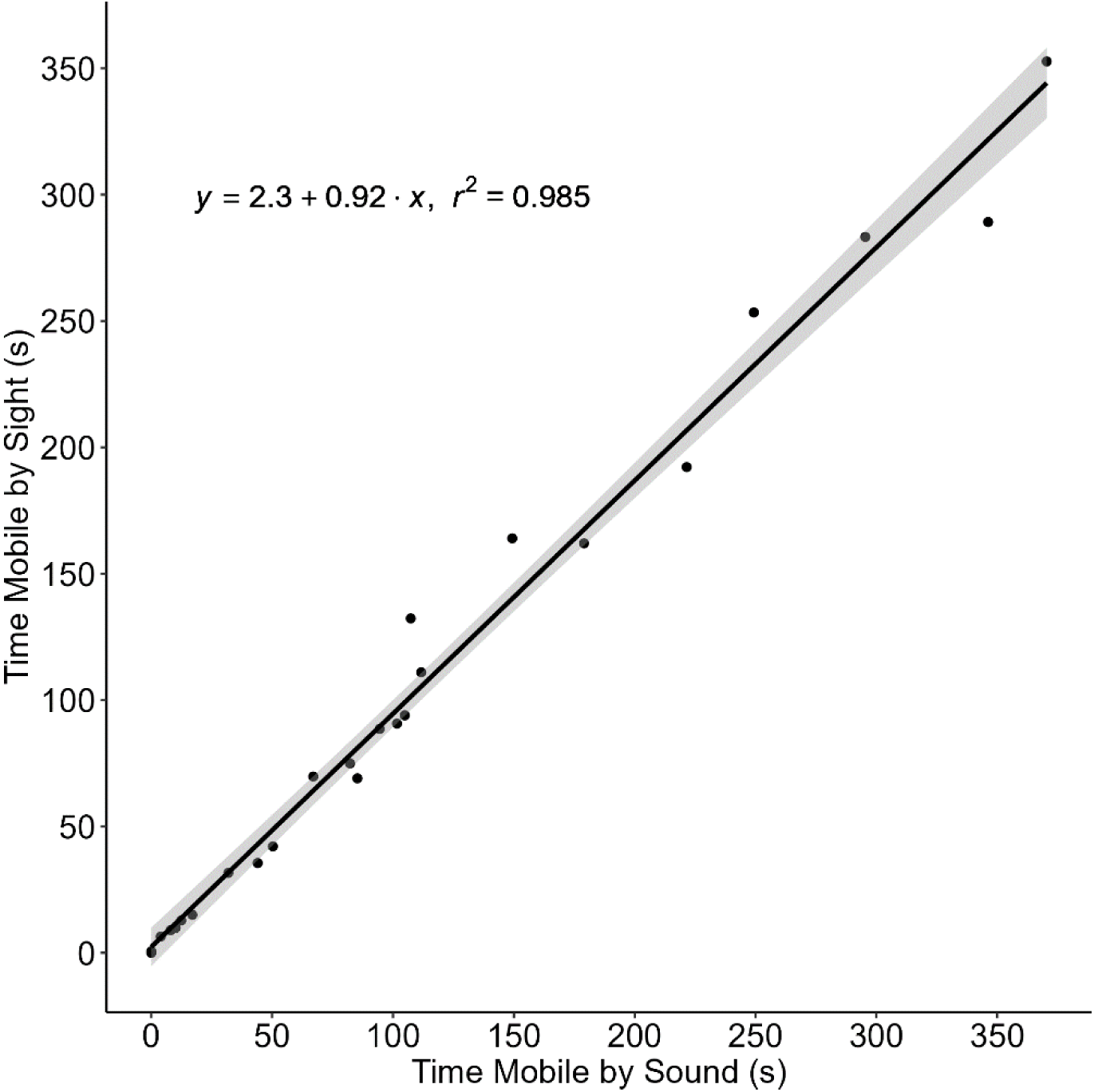
Comparison of time spent mobile by 25 pufflings in a 600 second trial in 25 lighted videos (including HPS, Warm White LED, and Cool White LED), as scored by watching the video (y-axis) versus listening to the video (x-axis). Note that there are two overlapping points around (0, 0).

